# Hexose translocation mediated by *Sl*SWEET5b is required for pollen maturation in *Solanum lycopersicum*

**DOI:** 10.1101/2021.08.30.458256

**Authors:** Han-Yu Ko, Hsuan-Wei Tseng, Li-Hsuan Ho, Lu Wang, Tzu-Fang Chang, Annie Lin, Yong-Ling Ruan, H. Ekkehard Neuhaus, Woei-Jiun Guo

**Affiliations:** Department of Biotechnology and Bioindustry Sciences, National Cheng Kung University, NO.1, University Road, Tainan City, Taiwan 7013; Plant Physiology, University of Kaiserslautern, Erwin-Schrödinger-Str., 22 D-67663, Kaiserslautern, Germany; School of Environmental and Life Sciences and Australia-China Research Centre for Crop Science, The University of Newcastle, Callaghan, New South Wales 2308, Australia

**Keywords:** Micro-tom, sugar unloading, *Solanum lycopersicum*, pollen maturation, SWEET, facilitator, male sterility

## Abstract

Pollen fertility is critical for successful fertilization and, accordingly, for crop yield. While sugar unloading affects growth and development of all types of sink organs, the molecular nature for sugar import to tomato pollen is poorly understood. However, SWEET transporters have been proposed to function in pollen development. Here, qRT-PCR revealed that *SlSWEET5b* was markedly expressed in flowers when compared to the remaining tomato *SlSWEET*s; particularly, in the stamens of maturing flower buds undergoing mitosis. Distinct accumulation of *Sl*SWEET5b-GUS fusion proteins was present in mature flower buds, especially in anther vascular and inner cells, symplasmic isolated pollen cells and styles. The demonstration that GFP fusion proteins located to the plasma membrane support the idea that the *Sl*SWEET5b carrier functions in apoplasmic sugar translocation during pollen maturation. Such function is in line with data from yeast complementation experiments and radiotracer uptakes, showing that *Sl*SWEET5b operates as a low affinity hexose-specific passive facilitator, with a K_M_ of ~36 mM. Most importantly, RNAi-mediated suppression of *SlSWEET5b* expression resulted in shrunken nucleus-less pollen cells, impaired germination and low seed yield. Interestingly, stamens from *SlSWEET5b*-silenced tomato mutants contained significantly lower amounts of sucrose and increased invertase activity, pointing to reduced carbon supply and perturbed sucrose homeostasis in this tissue. Taken together, our findings reveal an essential role of *Sl*SWEET5b in mediating apoplasmic hexose import into phloem unloading cells and into developing pollen cells to support pollen mitosis and maturation in tomato flowers.

**One-sentence Summary:** Plasma-membrane-localized SlSWEET5b facilitates a sequential hexose flux, from phloem to anther cells and from anther locule to pollen, to support pollen maturation and fertility in tomato flowers.

---

Pollen development is essential to generate the male gametophyte that is critical for plant reproduction and crop yield (Mascarenhas, 1989; Ma, 2005). In flowering plants, pollen cell development takes place in stamen, comprising anthers and filaments attached to the flower receptacle. Stamen development is characterized by active growth and a remarkable metabolic activity (Borghi and Fernie, 2017), and thus exhibiting high sink strength for nutrients during early stages of flower formation (Clément et al., 1996; Imlau et al., 1999). Due to limited photosynthesis in green sepals and young petals (Clément et al., 1997; Aschan et al., 2005), stamen and pollen growth depend on the import of carbon resources, predominately sucrose (Suc) (Ho and Nichols, 1975; Borghi and Fernie, 2017), from source leaves (Goetz et al., 2001; Müller et al., 2010). However, during pollen development, programmed degradation from surrounding maternal anther cells results in a complete symplasmic isolation of the male gametophyte (Scott et al., 1991; Clément and Audran, 1995). Accordingly, apoplasmic sugar transport mediated by plasma membrane localized carriers fulfils an essential role in acquiring carbon for pollen development (Truernit et al., 1999; Borghi and Fernie, 2017).

Generally, Suc is first unloaded from flower terminal phloem and allocated symplasmically to anther cells through the filament (Borghi and Fernie, 2017), representing a cluster of vascular tissues that are continuously connected to maternal flower phloem tissues in the receptacle (Mascarenhas, 1989). The symplasmic transfer into anther cells has been demonstrated by post-phloem transport of green fluorescing proteins in Arabidopsis anthers (Imlau et al., 1999). Subsequent to this, inner pollen cells receive their nutrients via apoplasmic transport across the boundary between tapetum and pollen cells. During early growth stage (from microspore mother cell to sporal mitosis), the tapetum, a nutritious cell layer that enclose the whole pollen sac and acts as a sugar buffer (Goldberg et al., 1993; Castro and Clément, 2007), forms an apoplasmic barrier between anther inner cells and pollen (Clément and Audran, 1995; Borghi and Fernie, 2017). Consequently, Suc must be exported from anther inner cells and imported into tapetum cells by sucrose transporters (SUT/SUCs) prior to apoplasmic release to the locule, representing the apoplasmic space around pollen cells (Scott et al., 1991).

In some species, such as tomato (*Solanum lycopersicum*), a large portion of the unloaded Suc can be hydrolyzed in filament and anther inner cells by cell wall invertases (CWINs), leading to glucose (Glc) and fructose (Fru) (Pressman et al., 2012). Accordingly, intercellular transport of these sugars has to be mediated by hexose importers (Borghi and Fernie, 2017). During the pollen maturation stage, tapetum cells are programed for degradation and thus the adjunct anther inner cells became the major nutritive reservoir to supply sugars apoplasmically for pollen development (Goldberg et al., 1993; Clément and Audran, 1995; Castro and Clément, 2007). In all cases, once sugars are unloaded into the locule, apoplasmically isolated pollen cells will take up either sucrose or hexoses by respective membrane transporters (Borghi and Fernie, 2017).

While many pollen-specific plasma membrane-localized SUT/SUC or STP type symporters have been characterized in various species (Ylstra et al., 1998; Truernit et al., 1999; Cheng et al., 2015; Borghi and Fernie, 2017), the major carriers responsible for apoplasmic sugar transfer for pollen development are not fully identified. Decreased activity of sucrose symporters, such as SUT1 in rice (Hirose et al., 2010), SUT2 in tomato (Hackel et al., 2006) or SUC1 in Arabidopsis (Sivitz et al., 2008) only affects pollen germination and pollen tube growth, but not pollen development. Down-regulation of the pollen-expressed sucrose transporter *Cs*SUT1 in male cucumber flowers led to reduced carbohydrate content and shriveled pollen grains, suggesting that *Cs*SUT1mediates sucrose import into cucumber pollen for their development (Sun et al., 2019).

By contrast, the molecular nature of major hexose transporters likely required for pollen formation - as discussed above - has yet to be determined. Several genes encoding hexose transporters exhibit significant expression in pollen cells (Truernit et al., 1999; Schneidereit et al., 2003; Büttner, 2010; Borghi and Fernie, 2017). Yet, no differences in pollen morphology were observed when expression of hexose transporters, such as cucumber *Cs*HT1 (Cheng et al., 2015), Arabidopsis *At*STP6 (Scholz-Starke et al., 2003) or *Petunia hybrida Ph*PMT1 (Garrido et al., 2006) was suppressed. However, the high hexose concentrations in pollen locule favor the involvement of energy-efficient facilitators, e.g. members of the large SWEET (sugar will eventually be exported transporters) facilitator family have been proposed as candidates to mediate hexose import into pollen cells.

Since the first description of the SWEET protein family (Chen et al., 2010), a growing body of evidence indicates that SWEET members are key players in sugar allocation between source and sink organs (Sauer, 2007; Chen, 2013; Eom et al., 2015; Julius et al., 2017). In general, based on amino acid similarity four clades of SWEET genes can be identified in various plant species (Chen, 2013; Eom et al., 2015). Particularly, clades I and II SWEET members exhibit high transport activity for hexoses, while clade III members mainly transport sucrose and some clade IV SWEET members locate intracellular transport of fructose across the vacuolar membrane. In source leaves, SWEET11 and 12 catalyze passive sucrose export from phloem parenchyma cells into the phloem apoplasm (Chen et al., 2012), prior to active sucrose loading into the phloem mediated by proton-coupled SUC2 (Bezrutczyk et al., 2018). In sink organs, different sets of SWEET facilitators cooperate with energy-coupled sugar carriers to mediate intercellular export or import of sucrose or hexoses, to accomplish the apoplasmic sugar unloading for development of sink organs, such as clade III SWEETs for seeds (Chen et al., 2015c; Bezrutczyk et al., 2018; Yang et al., 2018; Shen et al.; Wang et al., 2019), SWEET9 for nectarines (Lin et al., 2014), SWEET1 for new leaves (Ho et al., 2019), and SWEET15 for fruits (Ko et al., 2021).

In Arabidopsis, several SWEET encoding genes are expressed in pollen cells (Chen et al., 2015b). In particular, transcripts of *AtSWEET8* (*RPG1*) and *AtSWEET13* (*RPG2*) preferentially accumulate in male microspores and tapetum cells, since the development of microspore mother cells (Guan et al., 2008; Sun et al., 2013). Disruption of both *At*SWEET8 and *At*SWEET13 function via T-DNA insertion results in defective pollen wall formation and impaired pollen fertility (Guan et al., 2008), probably due to lack of sugars for synthesis of the pollen cell wall. Latter studies highlight the possibility that SWEET transporters may also participate in sugar translocation into pollen of crop species that are the center for food production. Considering diverse sugar contents in pollen cells across plant species, ranging from 20% to 60% in pollen’s dry mass (Conti et al., 2016), and various types of sugars used for intercellular translocation (Borghi and Fernie, 2017), further investigation is required to determine the role of SWEET members in pollen development on a crop-specific basis. For self-fertilized fruit crop species, such as tomato, the molecular mechanism underlying pollen formation might even become instrumental for directed breeding approaches aiming to generate male sterility for hybrid seed production (Du et al., 2020).

In this study, we discovered that tomato *SlSWEET5b*, belonging to clade II SWEETs, is specifically expressed in stamen of mature flowers. Examinations of GUS- and GFP-fusion proteins indicated that *Sl*SWEET5b functions in the plasma membrane of both, pollen and stigma cells. In conjunction with transport activity assays in yeast and analysis of transgenic knock-down tomato lines, we propose an essential role of *Sl*SWEET5b in hexose translocation into pollen grains, required for their maturation and fertility.

## Results

### *SlSWEET5b* mRNA accumulates specifically in tomato stamen

To determine if a specific member of the SWEET family is involved in sugar unloading during pollen development in tomato flowers, we performed quantitative RT-PCR for 30 tomato *SlSWEET* genes using cDNA isolated from whole flowers at 1 day post anthesis. *SlSWEET5b* transcripts were markedly more abundant in tomato flowers than all other *SlSWEET* genes (Fig. 1A) while this mRNA is only low abundant in all vegetative organs tested (Fig. 1B). When compared to closely related clade II *SWEET* homologs, only *SlSWEET5b* transcripts predominantly accumulated in stamens, but not in petals, sepals and ovary of flowers (Fig. 1C). Within the clade II *SlSWEET*s solely *SlSWEET5b* expression was also specifically induced during the late stage (S15, 7 mm) of flower development (Fig. 1D) (Brukhin et al., 2003). The close association between *SlSWEET5b* RNA expression and pollen maturation was not observed for other clade II SWEETs (Fig. 1C and D), suggesting that *SlSWEET5b* is likely involved in pollen maturation during flower development. Therefore, we focused on the functional characterization of *SlSWEET5b* thereafter.

**Figure 1.**
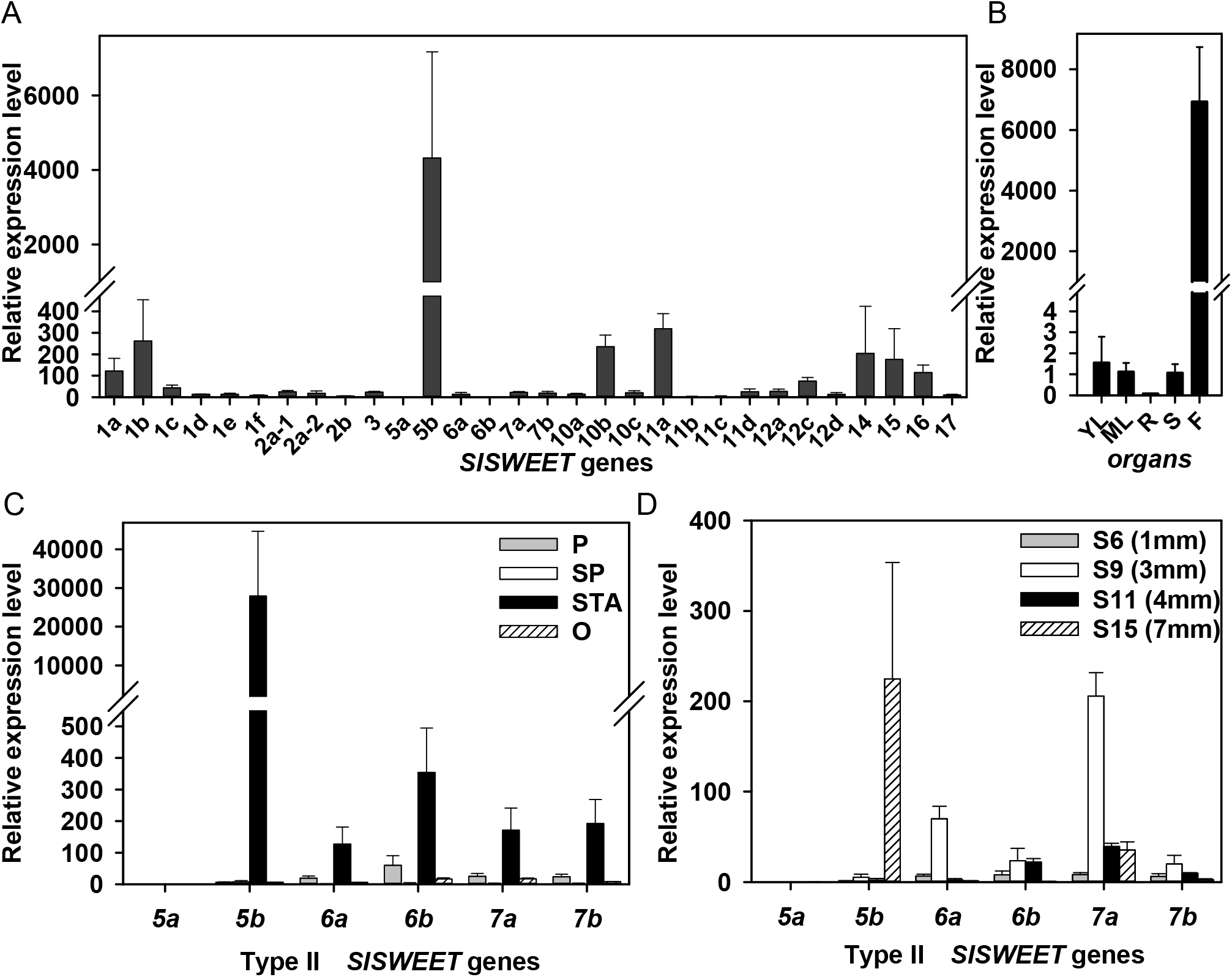
Dominant expression of *SlSWEET5b* in stamens of tomato flowers. A, Stable mRNA transcripts of 30 *SlSWEET*s in flowers. B, Developmental expression of *SlSWEET5b* in tomato. YL, young leaves. ML, mature leaves. R, roots. S, stems. F, flowers. C, Expression of clade II *SlSWEETs* in various flower organs. P, petal. SP, sepal. STA, stamen. O, ovary. D, Expression of clade II *SlSWEETs* during flower bud development. Stages (S) and sizes (mm) of flower buds were indicated. Total RNA was isolated from indicated organs from 6-week-old soil grown tomato plants and derived cDNA was used for RT-qPCR with gene-specific primers. Relative expression by normalizing to an internal control, *SlActin7*, is shown. Results are means ± SE from three independent biological replicates.

### *Sl*SWEET5b proteins accumulate in unloading cells and pollen

Several studies have shown that *Sl*SWEET protein abundance can be uncoupled to corresponding mRNA transcripts (Guo et al., 2014; Chen et al., 2015a). To investigate the tissue-specific expression of *Sl*SWEET5b proteins, we created transgenic tomato plants expressing a *Sl*SWEET5b-*β*-glucuronidase (GUS) fusion protein. The transgene included the complete genomic *SlSWEET5b* DNA sequence, containing all introns driven by the native *SlSWEET5b* promoter with the GUS coding sequence added as a C-terminal fusion. In T0 transgenic tomato plants, prominent blue staining, for GUS activity, was only observed in mature flowers, but not in roots, leaves, or young flower buds (Fig. 2A-D). In flowers, strong expression of *Sl*SWEET5b-GUS fusion was detected in the top region of the style (Fig. 2E), pollens cells (Fig. 2F), and inner anther cells surrounding the pollen sac (open arrow head, Fig. 2F). During fruit development, distinct GUS activities were additionally present in vascular tissues of pericarp, columella and seeds (Fig. 2G, H). To address tissue specificity more closely, histochemical staining of stamen sections was performed. In both longitudinal and transverse sections of pistil and stamen, expression of *Sl*SWEET5b-GUS was clearly detected in the upper region of style (arrow head, Fig. 2I) and mature round pollen grains pollen (Fig. 2I, J, K). In anthers, marked blue staining was also observed in unloading cells around xylem tissues, including phloem and surrounding parenchyma cells (Fig. 2L). Slight blue staining was also observed in the inner anther cell layer around the pollen sac (open arrow head, Fig. 2K).

**Figure 2.**
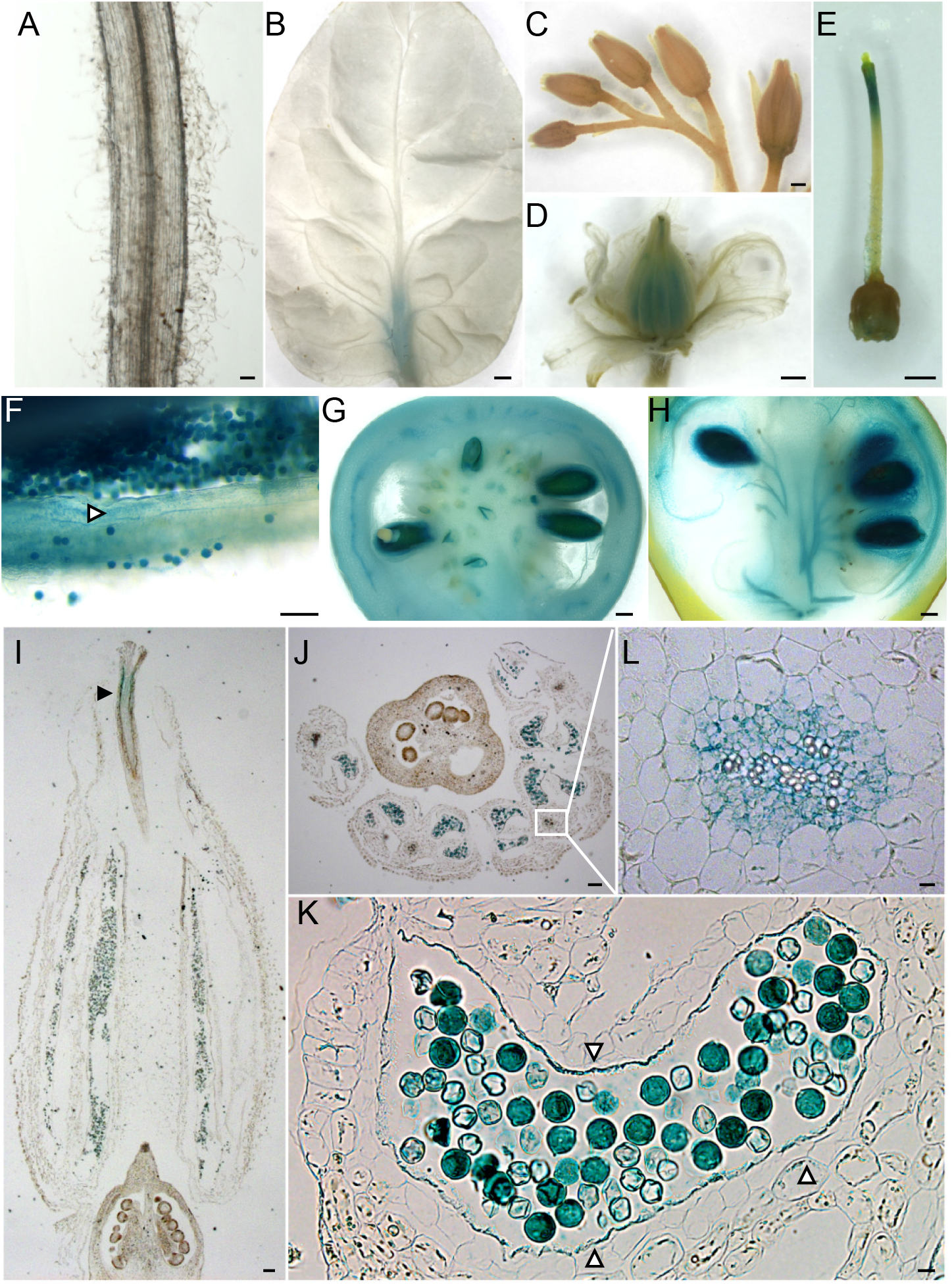
Specific accumulation of *Sl*SWEET5b in reproductive organs in tomato. Histochemical staining of GUS activities in transgenic tomato plants expressing the *Sl*SWEET5b–GUS fusion protein, driven by the native *SlSWEET5b* promoter. A, mature roots. B, mature leaflet. C, flower buds. D, open flower. E, pistil. F, inside of stamen. G, young green tomato fruit. H, mature red tomato fruits. I. Longitudinal section of a stamen and pistil. Arrow head indicate the stained style. J. Transverse section of a stamen. K, zoom-in picture of J. L, zoom-in picture of the white box in J. Open arrow heads in F and K indicate the inner cell layer around the pollen sac. Bars = 100 μm in A, I, J; 1 mm in B-H; 10 μm in K, L.

### Plasma-membrane localization of *Sl*SWEET5b

To examine the subcellular localization of *Sl*SWEET5b, we transiently expressed *Sl*SWEET5b-YFP fusion proteins driven by the 35S promoter in Arabidopsis protoplasts. The yellow fluorescence derived from *Sl*SWEET5b-YFP fusions was mainly located on the plasma membrane, which enclosing cytosolic chloroplasts, as indicated by their red autofluorescence (arrow head, Fig. 3). When co-transformed with the plasma membrane marker, *At*PIP2A-CFP, bright fluorescence resulting from overlapping yellow and blue signals was observed on the plasma membrane (Fig. 3), demonstrating that *Sl*SWEET5b located primarily on the plasma-membrane *in planta*.

**Figure 3.**
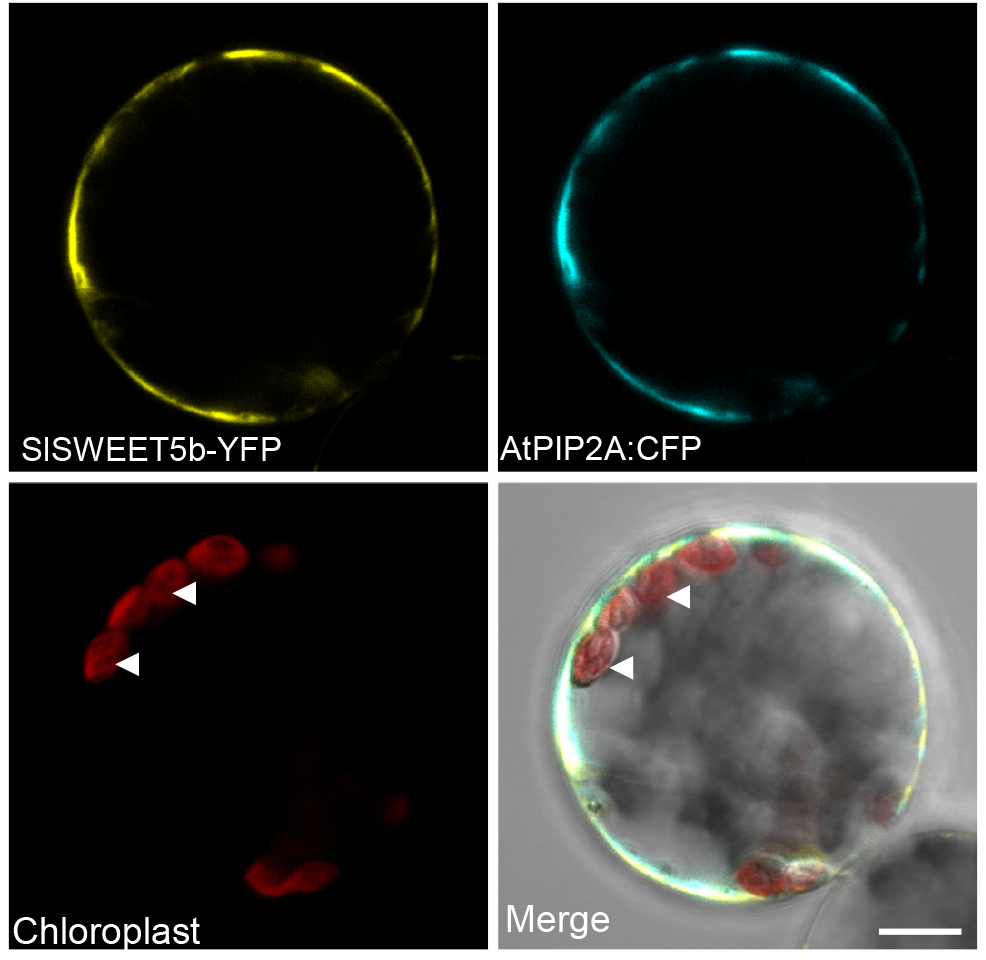
Subcellular localization of *Sl*SWEET5b on the plasma membrane. Arabidopsis protoplast was coexpressed with the *Sl*SWEET5b-YFP protein and the plasma membrane marker, *At*PIP2A-CFP. All YFP and CFP fluorescence, auto-fluorescence from chloroplasts (arrow heads), and merged signals from the same focal plane is shown. Bar = 10 μm.

### Hexose-specific transport activity of *Sl*SWEET5b in yeast

*Sl*SWEET5 belong to clade II SWEET proteins, and these proteins generally catalyze transport of hexoses, as exemplified by Arabidopsis *At*SWEET5 (Chen et al., 2010). To verify sugar transport properties of *SI*SWEET15b, we expressed the gene in the baker’s yeast mutant EBY4000, which lacks import activities for hexoses and does not grow on hexose-containing media (Wieczorke et al., 1999). As expected, expression of the control yeast hexose transporter *Sc*HXT5 enables growth on medium containing 2% Glc or Fru (Fig. 4A). Neither the empty vector cells nor the sucrose carrier *At*SUC2 were able to restore growth of the yeast mutant. Similar to *Sc*HXT5, expression of *Sl*SWEET5b complemented the growth of EBY4000 mutant cells on both hexose-containing media (Fig. 4A). Analysis of [^14^C]-glucose ([^14^C]Glc) uptake confirmed that yeast cells expressing *Sl*SWEET5b imported Glc in a time dependent manner, whereas cells containing the empty vector accumulated much less Glc (Fig. 4B). The kinetic analysis of the Glc transport activity of *Sl*SWEET5b revealed a K_M_ value of 36 mM for Glc (Fig. 4C)

**Figure 4.**
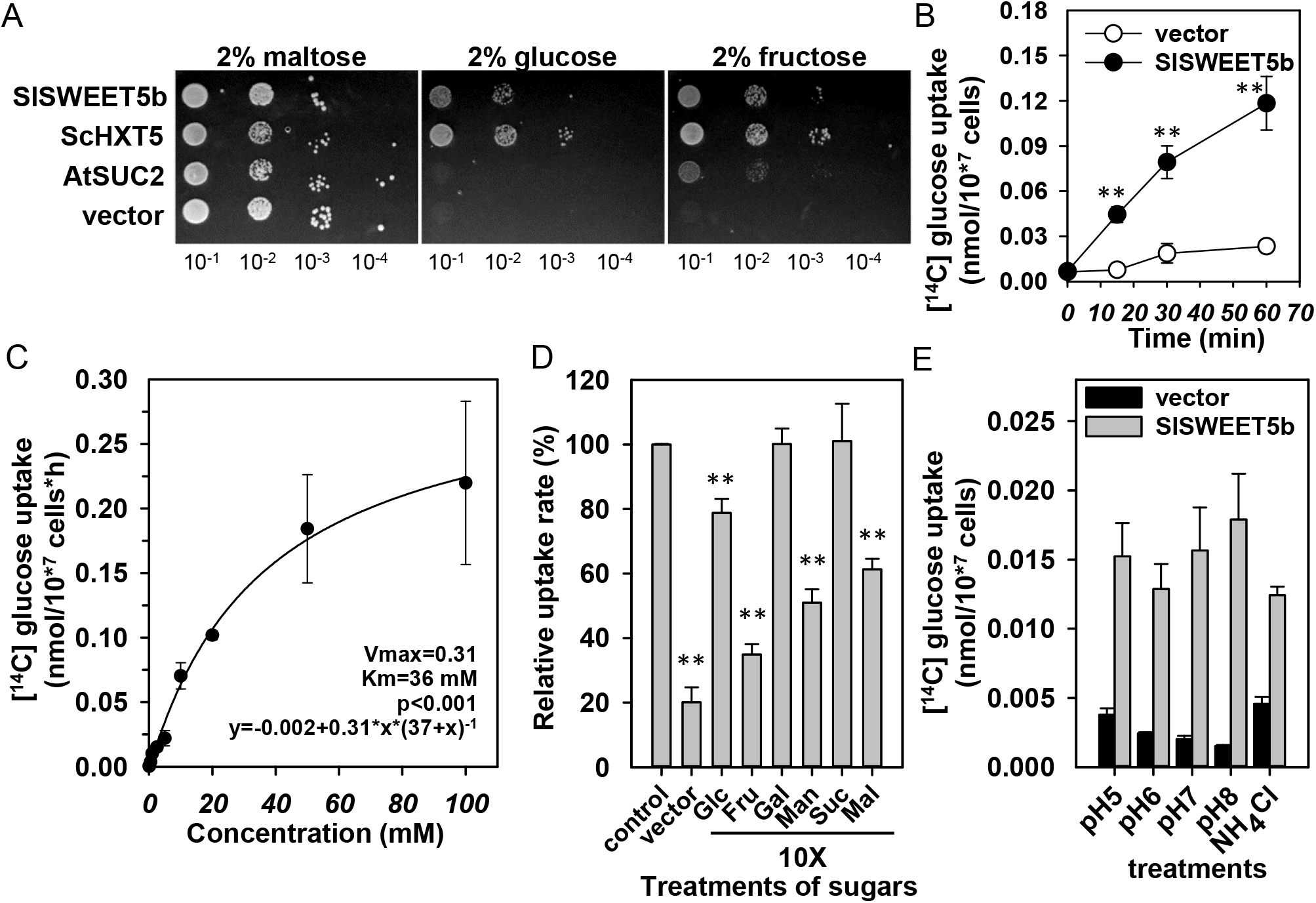
Transport activities of *Sl*SWEET5b to hexoses in yeast. A, Complementation of hexose import by *Sl*SWEET5b in the yeast mutant. Yeast transformants expressing the yeast hexose transporter (*Sc*HXT5), Arabidopsis sucrose symporter (*At*SUC2) or the empty vector (vector) were serially diluted (10-fold) and cultured on solid media supplemented with 2% maltose, glucose, or fructose, respectively. Yeast growth was imaged after incubation at 30°C for 4 to 6 d. B, Time-dependent [^14^C]Glc uptake activity. C, Kinetic of [^14^C]Glc uptake catalyzed by *Sl*SWEET5b. D, Substrate specificity of *Sl*SWEET5b to hexoses. Relative uptake rates of various sugars was measured by incubating cells expressing *Sl*SWEET5b with [^14^C]Glc only (control) or with 10 times concentrated sugars (Glc, glucose; Gal, galactose; Fru, fructose; Man, mannose; Suc, sucrose; Mal, maltose). The uptake rate of cells expressing the empty vector (vector) was also shown. E, Insensitivity of *Sl*SWEET5b transport to alkaline pH. Yeast cells expressing the empty vector or *Sl*SWEET5b were subjected to various pH conditions and treatment of NH_4_Cl. Results of [^14^C]Glc uptake are shown. B-E, Results are mean ± SE (n = 3 to 4 independent colonies). Significant differences from the empty vector (B), the control (D) or the pH 5 condition (E) were determined by Student’s t-test, respectively. **: P< 0.01.

To address the substrate specificity of *Sl*SWEET5b, a 10-fold excess of unlabeled sugars was added to the medium to compete with [^14^C]Glc. Relative uptake rates showed that only Glc, Fru, and mannose, but not galactose or sucrose, could compete the binding and significantly reduce import of [^14^C]Glc (Fig 4D). The decreased Glc uptake in the presence of maltose is probably due to background activity in EBY4000 yeast strain, as shown previously (Chen et al., 2010; Ho et al., 2019). Moreover, the Glc transport catalyzed by *Sl*SWEET5b was very similar under different pH ranging from, acid (pH 5) to alkaline (pH 8), and decreased only slightly upon the addition of the protonophore NH_4_Cl (10 mM) (Fig. 4E) and is probably due to cell toxicity of this compound (Ho et al., 2019). In summary, these results indicate that *Sl*SWEET5b is a proton-independent passive hexose facilitator with low substrate affinity.

### Loss of *SlSWEET5b* expression impeded seed production

To examine the function of *Sl*SWEET5b during pollen development, we generated transgenic knock-down tomato mutants expressing a RNA interference (RNAi) construct targeted to *SlSWEET5b* (*SlSWEET5b-RNAi*). In 20 independent T0 transgenic tomato plants (RNAi-1 to 20) the RNAi-6, 10, 16 lines showed significantly decreased levels of *SlSWEET5b* mRNA, ranging from 10% to 40% of the expression level present in the empty vector controls (Supplemental Fig. S1A). When compared to non-silenced (NS) T0 transgenic plants (lines RNAi-7, 11, 18, Supplemental Fig. S1A), no obvious differences in plant size and height were observed (Supplemental Fig. S1B). However, the severely silenced T0 line RNAi-6 exhibited strongly decreased total fruit number and fruit fresh weight. These changes were not observed in RNAi-10 and −16, lines with moderate reduction of RNA expression (Supplemental Fig. S2A, B). There were no consistent differences in average fruit yield and fresh weights between silenced RNAi lines (combined data from 3 RNAi lines, RNAi-HS) and NS transgenic plants (Fig. 5A, B). However, total seed numbers in three silenced lines (RNAi-HS) were markedly reduced, when compared to NS lines (Supplemental Fig. S2C; Fig. 5C). Only one seed each was obtained from RNAi-6 and 16 lines (Supplemental Fig. S2C). These results clearly demonstrated that reduced expression of *SlSWEET5b* significantly inhibited seed production.

**Figure 5.**
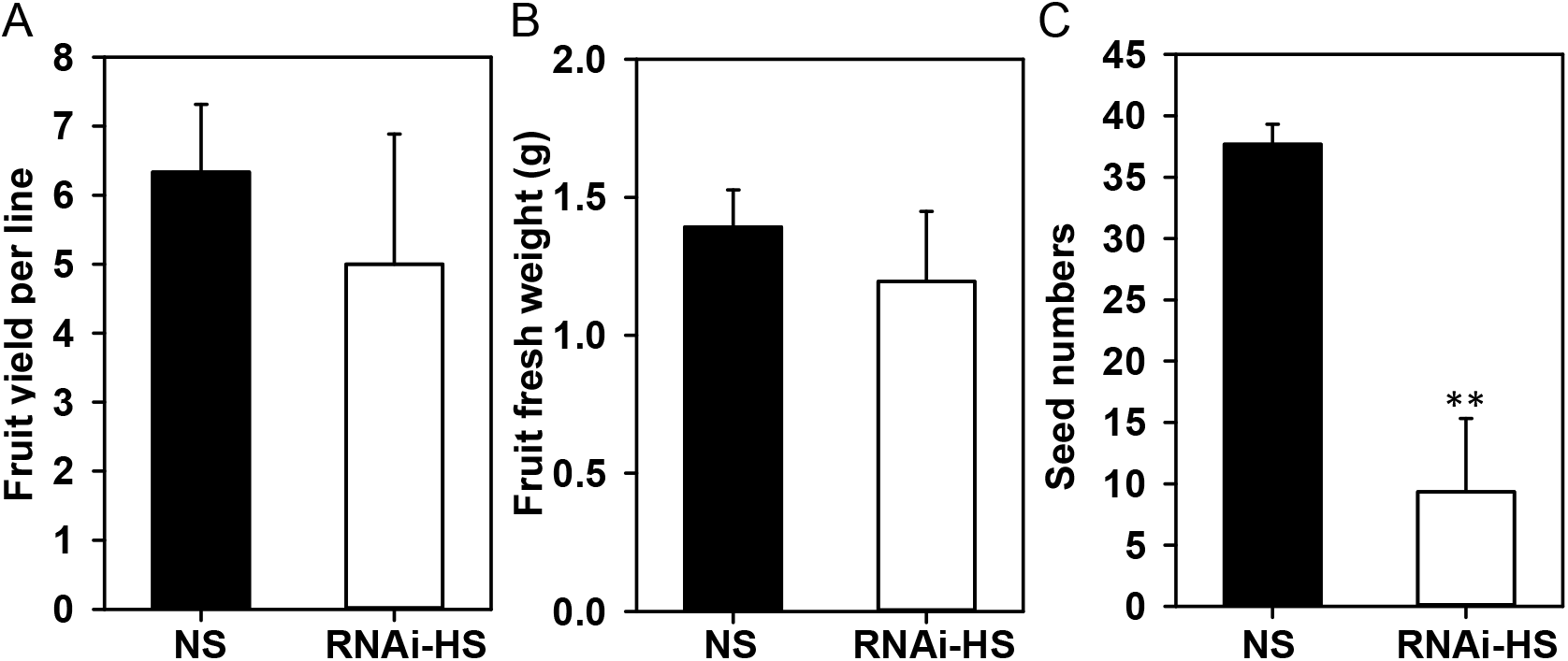
Fruit and seed yields of *SlSWEET5b* knock-down tomato plants. Mature red fruits were harvested from T0 transgenic tomato plants expressing the *SlSWEET5b-RNAi* construct. A, total fruit numbers. B, average fresh weights of each fruit. C, total seed numbers from each line. A to C, results are mean ± SE from 3 independent *SlSWEET5b*-silenced lines (RNAi-HS). Significant differences from non-silenced transgenic lines (NS) were analyzed by Student’s t-test. **: P<0.01.

### Silencing *SlSWEET5b* expression resulted in male sterility

Based on dominant localization of *Sl*SWEET5b proteins in pollen and unloading cells (Fig. 2J, L), we assumed that the decreased seed production in corresponding mutant plants may result from pollen fertility. The lack of fertile T1 seeds from both RNAi-6 and RNAi-16 lines prevented further characterization of these lines (Supplemental Fig. S2C). Therefore, we examined pollen development in T1 plants derived from the moderate silenced RNAi-10 line. Fifteen T1 plants were assayed by RT-qPCR and three lines, RNAi-10-2, 10-8, and 10-16, exhibited distinct silencing of *SlSWEET5b* gene expression in flowers to approximately 10% of WT flowers (Supplemental Fig. S3A). Thus, these highly silenced T1 lines (T1-HS) were characterized in more details. Similar to T0 plants, no significant differences in plant size (Supplemental Fig. S3B) and average fruit fresh weights (Supplemental Fig. S3C) were observed between WT, non-silenced (T1-NS) and T1-HS plants. However, the germination rate of pollen grains from T1-HS plants was only ~5 %, compared with 40% for WT pollen (Fig. 6A). Staining with the DNA-binding dye DAPI solution (4’,6-diamidino-2-phenylindole, Vector Laboratories Inc.) demonstrated the presence of vegetative and sperm nuclei in WT pollen (Fig. 6B). In contrast, nuclei were absent in most T1-HS pollen and they were also misshapen (Fig. 6B). Up to 80% of *SlSWEET5b*-silenced pollen grains were of abnormal shape and lacked nuclei, compared to less than 25% in WT flowers (Fig. 6C). A cryo-electron microscopic analysis revealed that pollen cells from *SlSWEET5b*-silenced plants (T1-HS) tended to clump, stayed adhered to tapetum cells, with a shrunken, invaginated shape. In contrast, WT pollen appeared of smooth and round shape and were freely released into the anther locule (Fig. 6D).

**Figure 6.**
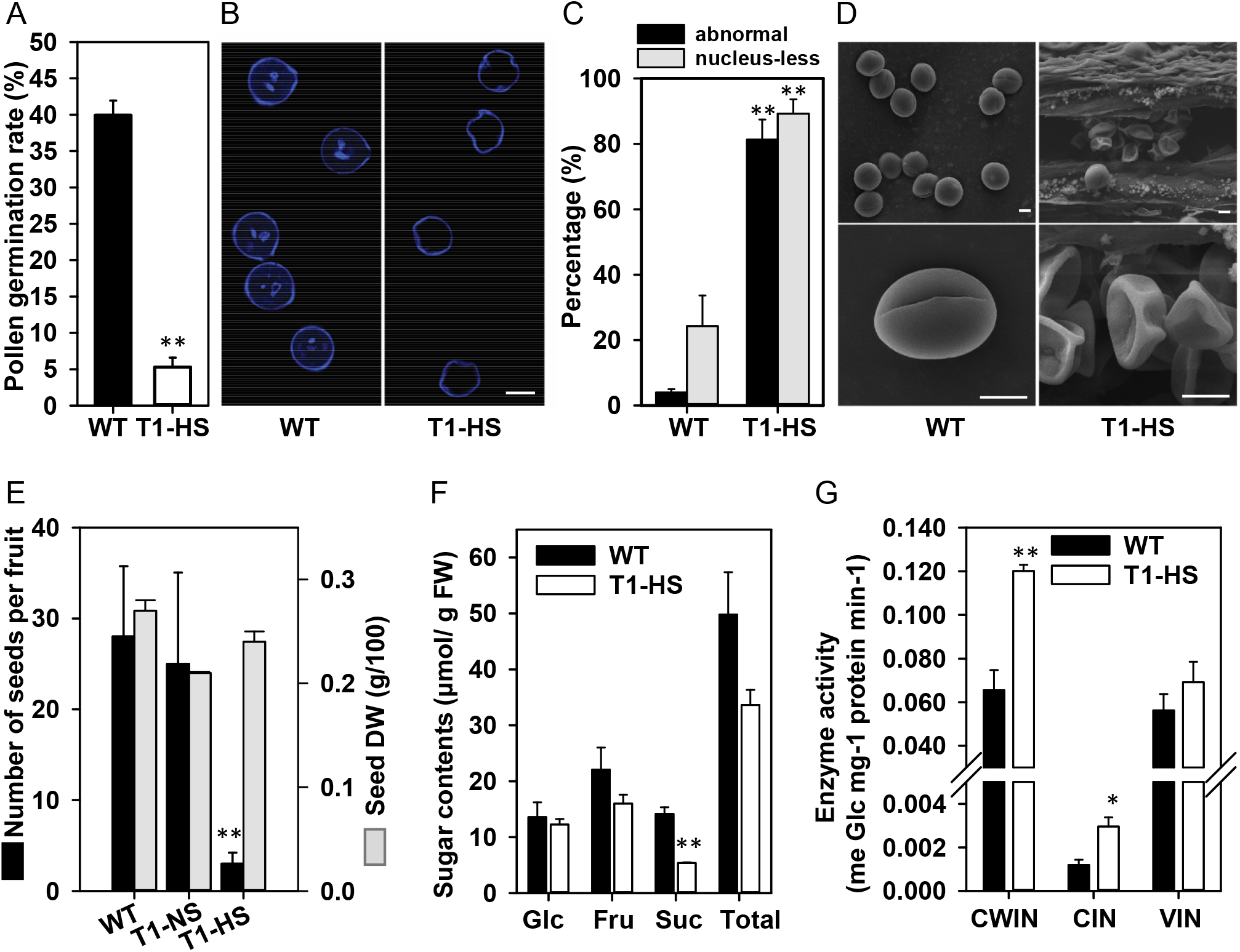
Aborted pollen and seed development in *SlSWEET5b*-silenced tomato plants. A, pollen germination rate. Germinated pollen of wild type (WT) and T1 highly silenced plants (T1-HS) were counted after incubation in germinating buffer for 4 h. B, pollen morphology in (A). Pollen grains freshly harvested from tomato flowers were stained with the DAPI stain for 30 min and imaged under a confocal microscope. C, ratio of defective pollen grains in (B). D, electron microscopic pictures of fresh pollen in tomato stamens. Stamens were detached from 1-d-anthesis flowers of WT and T1-HS plants and directly frozen on the slide for observation. E, seed yields of T1 transgenic tomato plants. Results are mean ± SE from 5 independent fruits that were harvested from WT, T1-HS or transgenic plants without silencing of *SlSWEET5b* expression (T1-HS). F, sugar contents of stamen. G, invertase activities of flowers. In F and G, Stamens or flowers were harvested from 1-d-anthesis flowers of WT and T1-HS lines. Sugar contents (Glc, glucose; Fru, fructose; Suc, sucrose; Total, sum of three sugars) and invertase activities (CWIN, cell wall invertase; CIN, cytosolic invertase; VIN, vacuolar invertase) were analyzed. Results in A, C, F, G are mean ± SE from 3 to 4 biological samples collected from independent flowers. Bar = 10 μm in B and D. Differences from WT were analyzed by Student’s t-test. *, **: P<0.05, 0.01.

Consistent with defective pollen, seed number per fruit was strongly decreased by 90 %, in T1-HS mutants compared to WT or T1-NS plants (Fig. 6E). The small number of T2 seeds that did develop in T1 *SlSWEET5b*-silenced plants (T1-HS) were subsequently shown to be non-transgenic segregants derived from the heterozygous T1 plants. Dry weight of T2 seeds from T1-HS plants was similar to those of WT and T1-NS plants (Fig. 6E), indicating that SlSWEET5b does not participate in seed filling upon fertilization.

### Sugar homeostasis was affected in *Sl*SWEET5b-silenced stamens

Based on the hexose transport activity of *Sl*SWEET5b in yeast (Fig. 4), we hypothesized that the defective pollen observed in silenced plants may be due to sugar starvation. To examine this hypothesis, we compared sugar contents in stamens of open flowers of T1-HS lines. Significant reductions of sucrose were observed in stamens of silenced plants (Fig. 6F). Lower fructose and total sugar contents were also observed in T1-HS stamens. Studies have showed that hexose levels in pollen are regulated by extracellular invertase activity (Shen et al., 2019; Goetz et al., 2001). Significantly higher activities of cell wall invertase (CWIN) and cytosolic invertase (CIN) were measured in the silenced flowers (T1-HS), compared to WT (Fig. 6G). However, there was no difference in vacuolar invertase activities (VIN). In sum, latter results confirm that silencing *Sl*SWEET5b transport activity results in reduced sugar accumulation in stamen and perturbed sugar metabolism by altering invertase activity.

## Discussion

It is generally accepted that optimization of molecular properties to support higher fruit yield is of great importance for global food security (Ruan et al., 2012). Tomato (*Solanum lycopersicum*) is one of the major fruit crops showing a global production value up to 90 million US$ (FOASTAT, 2018). Fruit yield in tomato has been shown to be closely related to pollen fertility (Xu et al., 2017), which critically depends on continuous supply of carbohydrates, especially sugars (Clément et al., 1996; Firon et al., 2006; Castro and Clément, 2007).

During pollen maturation, high amounts of sugars are translocated to tomato stamen to support cell wall formation and pollen germination subsequent to pollination (Pressman et al., 2012). However, symplastic isolation of developing pollen cells from surrounding nutritive anther cells, such as tapetum cells (at early growth stage) or the anther inner cells (during pollen maturation stage; Polowick and Sawhney, 1993; Brukhin et al., 2003), results in the need for a set of plasma-membrane sugar transporters to mediate two steps of sugar flux. During a first transport event, sugars are exported from anther cells into the locular apoplasm and a second flux provides sugars into developing pollen (Clément and Audran, 1995; Borghi and Fernie, 2017). Results presented here indicate that *Sl*SWEET5b is the primary sugar carrier responsible for hexose translocation for pollen maturation in tomato flowers.

Among 30 *SlSWEET* genes, *SlSWEET5b* was the sole member that show a dominant expression in flowers, while its expression in vegetative tissues is relatively low (Fig. 1A, B). The predominant expression of *SlSWEET5b* in stamen (Fig. 1C) was similar to the expression of previously characterized sugar carriers that function in sugar transport during pollen development, such as Arabidopsis *At*SWEET8 and *At*SWEET13 (Guan et al., 2008; Sun et al., 2013), and cucumber *Cs*SUT1 (Sun et al., 2019). Obviously, this positive correlation suggests a specific role for *Sl*SWEET5b in pollen development. In line with this we observed that the expression of *SlSWEET5b* was particularly high in the maturation stage of flower buds (S15, 7mm in Fig. 1D), when mitosis in the microspore and degradation of tapetum cells occur (Brukhin et al., 2003). In development stage 15 of a tomato flower, carbohydrates stored in tapetum cells are mobilized as soluble sugars to the locules (Goldberg et al., 1993; Castro and Clément, 2007). This coordinated process leads in both, tomato locule and pollen cells, to an accumulation of sugars gradually during maturation of microspores, reaching maximal levels prior to anthesis (Pressman et al., 2012).

The clear concurrence of a tissue specific *SlSWEET5b* expression in stamen with sugar accumulation in tomato pollen cells (Pressman et al., 2012) indicate that *Sl*SWEET5b is involved in sugar influx into pollen required for maturation. This assumption gains further supported by the accumulation of *Sl*SWEET5b protein in maturing pollen, but not in early developing miscrospores (Fig. 2C), as indicated by the analysis of GUS labelling using a whole-gene translational GUS fusion construct (Fig. 2). Moreover, the presence of *Sl*SWEET5b was also detected in corresponding sugar unloading cells, including the vascular tissue (Fig. 2L) and inner cell layers of the anthers (Fig. 2F, K). The anther phloem and the surrounding phloem parenchyma tissue represent the major sugar unloading sites for sucrose delivered from maternal filaments (Mascarenhas, 1989). When tapetum cells become degraded during pollen maturation, the anther inner cells, symplasmically isolated from pollen cells, serve as the main sugar reservoir for pollen development (Goldberg et al., 1993; Clément and Audran, 1995; Castro and Clément, 2007). Within stamen, the highest sugar concentrations are present in the anther cells compared to those in locule and pollen cells (Pressman et al., 2012). Thus, the localization of *Sl*SWEET5b in cells involved in apoplasmic sugar unloading in stamen supports the conclusion that *Sl*SWEET5b mediates the whole apoplasmic translocation pathway in stamen, initial transport into phloem parenchyma cells, sugar efflux from nutritive anther cells, and influx into pollens to support their development and maturation.

Latter assumption is further supported by independent observations: Firstly, *Sl*SWEET5b locates, similar to other tomato SWEET proteins involved in apoplasmic sugar exchange (Ho et al., 2019; Ko et al., 2021), in the plasma membrane (Fig. 3). Secondly, similar to clade II SWEET transporters in other species (Chen et al., 2010; Chong et al., 2014; Sosso et al., 2015), *Sl*SWEET5b transports Glc and Fru at comparable low affinities (Fig. 4A-D). Latter fact fits with high monosaccharide contents (10-fold higher than sucrose) in tomato anther cells and locule (and other *Solanacea* species) (Pressman et al., 2012). Thirdly, *Sl*SWEET5b acts as a facilitator (Fig. 4E) and as that represents a passive and bi-directional hexose-specific (Fig. 4A-D) carrier (Chen et al., 2010; Lin et al., 2014; Ho et al., 2019), which supports its role in dual transport during sugar unloading and loading during pollen development. The passive transport nature of *Sl*SWEET5ba concurs with physiological sugar gradients within the pollen sac, where glucose and fructose comprise 80% of total soluble sugars in the locular fluid, while these monosaccharides contribute to only 5% of total soluble sugars in pollen cells (Pressman et al., 2012). Thus, an approximate 10-fold concentration gradient of hexoses across locule fluid and the pollen cytoplasm allows for net uptake into the pollen cells, facilitated by *Sl*SWEET5b (Fig. 7).

**Figure 7.**
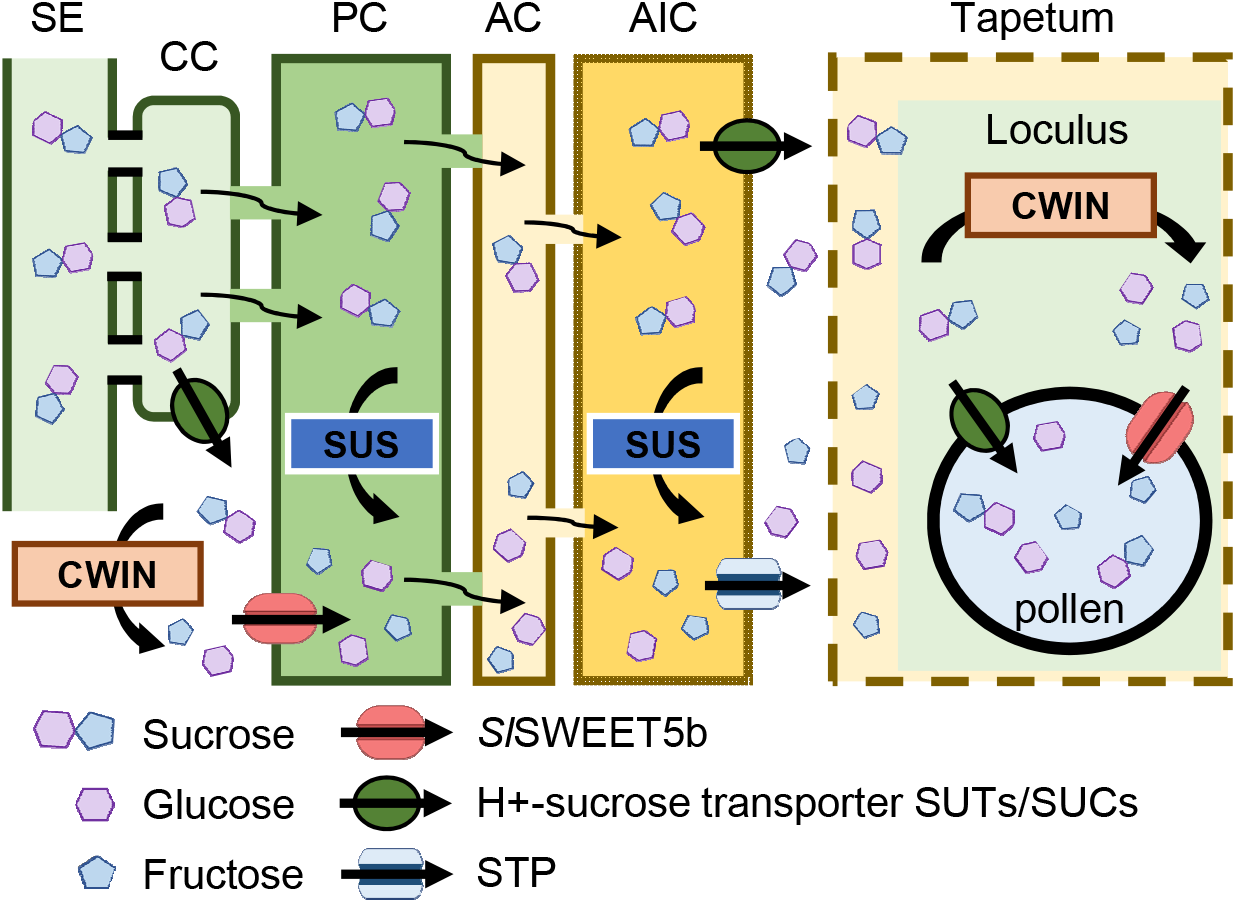
The functional model of *Sl*SWEET5b during pollen maturation. During pollen maturation, sucrose is translocated symplasmically to anther vascular tissues (SE, sieve element; CC, companion cell), where parts of sucrose are exported to vascular apoplasm in tomato stamens. Extracellular sucrose is then hydrolyzed by cell wall invertase (CWIN) to glucose and fructose. In this case, plasma membrane-localized *Sl*SWEET5b mediates import and export of apoplasmic hexoses to facilitate intercellular sugar distribution between phloem parenchyma cells (PC) and anther cells (AC). Due to symplasmic discontinuity between anther inner cell (AIC) and degrading tapetum cells, STP hexose transporter and some sucrose transporter may catalyze hexoses and sucrose fluxes into anther loculus space, where most sucrose would be hydrolyzed to hexoses by CWIN. Subsequent to this, *Sl*SWEET5b on the plasma membrane is required to efficiently import hexoses from the apoplasmic loculus into developing pollen for maturation and wall formation.

Interestingly, *Sl*SWEET5b proteins were also detected in styles (Fig. 2I). Our observation is fully consistent with stimulated expression of the *SlSWEET5b* gene in the style in response to pollination (Shen et al., 2019). Upon pollination, the pollen tube performs a rapid tip growth to deliver the sperm nuclei to ovules for fertilization (Lord and Russell, 2002). The high energy requirement for pollen tube growth depends on a continuous supply with nutrients in form of amino acids and sugars (Konar and Linskens, 1966; Borghi and Fernie, 2017), secreted by the surrounding style tissues (Mascarenhas, 1993; Cheung, 1996; Goetz et al., 2017). However, an apoplasmic barrier with the surrounding style tissues (Scott et al., 1991) predicts that plasma membrane-localized sugar transporters are required to allow transport of sugars, from style tissue to apoplasm and, subsequent to this, into elongating pollen tubes (Reinders, 2016; Goetz et al., 2017). We assume that *SlSWEET5b* may contribute to sugar unloading from style cells into the transmitting track, where sugar uptake into growing pollen tubes is mediated by proton driven SUC/SUT and HT type transporters (Cheng et al., 2015; Shen et al., 2019; Li et al., 2020).

The involvement of *Sl*SWEET5b in pollen development is further indicated by analyses on tomato mutants exhibiting decreased *SlSWEET5b* gene expression, as latter plants exhibited abnormal pollen morphology and induced male sterility (Fig. 6A-D). The compressed shape of *SlSWEET5b*-silenced pollen cells (Fig. 6D) is similar to those pollen in which cell wall formation is retarded due to disruption of the sugar transfer pathway (Guan et al., 2008; Sun et al., 2013; Sun et al., 2019).

It is surprising but explainable that *SlSWEET5b*-silenced plants contain decreased levels of sucrose in stamen tissue (Fig. 6F), although this carrier prefers monosaccharides and does not transport sucrose (Fig. 4C, D). It is known that sucrose is the major type of sugar present in mature pollen, although glucose and fructose are preferred carbon skeletons imported into tomato pollen (Pressman et al., 2012). However, it has been proposed that sucrose is constantly resynthesized in tomato pollen during maturation, most likely via fructokinase2 and sucrose-phosphate synthase activity (Pressman et al., 2012; Borghi and Fernie, 2017).

Such dynamic sucrose fluxes might act as the carbon buffer to support synthesis of pollen cell wall or accumulation of starch (Sun et al., 2013), latter functioning as an energy reserve for germination and representing a check point for pollen maturation (Datta et al., 2002). In maize, reduced levels of sugars in microspores correlate with starch deficiency and results in male sterility (Datta et al., 2002). In tomato, impaired *Sl*SWEET5b activity leads to impaired hexose import and carbohydrate deficiency, which limits both, cellular energy provision for mitosis and carbon precursor availability required for pollen maturation. Such processes ultimately result in male sterility and consistent with this view, viable seeds cannot be generated from pollen carrying severe *SlSWEET5b* silencing (Fig. 6E).

That decreased *Sl*SWEET5b activity negatively affect the pollen carbohydrate status is further indicated by the induced invertase activities, in particular the cell-wall invertase (CWIN), in flower tissue from *SlSWEET5b* silenced plants (Fig. 6G). High activity of CWIN was previously reported in tomato pollen (Pressman et al., 2012), where CWIN bound to the wall of pollen hydrolyzes Suc in the locular space to hexoses (Goetz et al., 2001; Hirsche et al., 2009). Similar to the effects observed in pollen from *SlSWEET5b*-silenced plants, decreased CWIN expression in tomato or tobacco leads to severe male sterility due to hexose starvation (Zhang et al., 2020). We propose that impaired monosaccharide import into pollen from *SlSWEET5b*-silenced plants is sensed as a low energy state and is genetically responded in a way leading to increased activity CWIN, which may generate sugar signals to stimulate sucrose hydrolysis or hexose uptake (Liao et al., 2020).

Taken together, we propose a functional model of *Sl*SWEET5b during pollen maturation (Fig. 7). During pollen maturation stage, a large amount of sugars are required for microspore mitosis and for formation of the structurally complex pollen cell wall. To fulfill the corresponding high carbon demand, in addition to symplasmic sucrose unloading, the plasma membrane located *Sl*SWEET5b protein facilitates sucrose unloading from phloem cells, by importing apoplasmic hexoses, previously hydrolyzed by CWIN from extracellular sucrose. Once sugars are released to the locule, most of the sucrose is converted to Glc and Fru by CWIN. The high monosaccharide concentration in the locule allows *Sl*SWEET5b to catalyze uptake of hexoses into maturing pollen cells. In tomato, pollen develops inside a closed anther cone, where individual stamen is fused to cover the whole pistil (Brukhin et al., 2003). Because of this morphology, self-fertilization is typical in tomato flowers driving the costs for hybrid seed production. Thus, induced male sterility by inactivation of *Sl*SWEET5b may provide a new strategy to develop male sterile lines for commercial F1 hybrid seed production (Du et al., 2020).

## Materials and Methods

### Plant materials and growth conditions

Tomato (*Solanum lycopersicum*) Micro-tom was used in this study. Seeds were sterilized, germinated and grown in a controlled environment room as described previously (Ho et al., 2019). For developmental expression profiling, roots, stems, young leaves (<2 cm long) mature leaves (>3 cm, terminal leaflet) and flowers (1 d post anthesis) were collected from 5-week-old hydroponically grown tomato plants. Flower organs (sepal, petal, stamen and ovary) and different stages of flowers buds (1, 3, 4, 7 mm length corresponding to stage 6, 9, 11, 15 developmental stage, respectively) (Brukhin et al., 2003) were harvested from 6- to 7-week-old soil grown tomato plants. All samples were stored at −80°C before analysis.

### RNA expression analysis

Total RNA was isolated from tomato organs with TRIsure reagent (Bioline, http://www.bioline.com/) along with an RNA purification column (GeneMark, http://www.genemarkbio.com/). The derived cDNA samples were used for RT-qPCR with gene-specific primers as described previously (Ho et al., 2019).

### Observations of GUS fusion proteins

The *SlSWEET5b* (Solyc06g071400) promoter (2000 bp upstream to ATG) was amplified from genomic DNA with specific primers (SWT5b-promoter-F and SWT5b-promoter-R, Supplemental Table S1) and cloned into the binary vector pUTKan via SacI and SacII sites (pUTKan-P_*SWEET5b*_). The *SlSWEET5b* genomic opening reading frame, including all introns (1110 bp after ATG) but lacking the stop codon, was amplified with primers (SWT5b-gDNA-F and SWT5b-gDNA-R, Supplemental Table S1) and cloned into the pUTKan-P_*SWEET5b*_ vector via SacII and BamHI sites. The resulting plasmid pUTKan-P_*SWEET5b*_::gSlSWEET5b was transferred into tomato plants by Argrobacterium transformation performed by the Transgenic Plant Core Lab in Academia Sinica (http://abrc.sinica.edu.tw/transplant/ index.php). T0 transgenic plants were regenerated from kanamycin resistant callus and grown in soil mix after shooting. T1 seeds were collected from mature red fruits.

Various organs harvested from T0 plants were stained histochemically for GUS activity and imaged after incubation of 16 h under a dissecting microscope as described previously (Ko et al., 2021). For tissue sections, flowers from 8-week-old T0 transgenic plants were collected and vacuumed-infiltrated with 2% PFA buffer (44 mM sodium hydroxide, 0.125% glutaraldehyde, 0.05% triton X-100, 0.05% Tween-20, 5 g paraformaldehyde, pH 7) for 10 min at 4 °C in the dark. Samples were then washed three times with PBS (Phosphate buffered saline) and stained for GUS activity for 16 h. Samples were washed with PBS and vacuumed-infiltrated with 4% PFA buffer (87.5 mM sodium hydroxide, 0.25% w/v glutaraldehyde, 0.1% v/v triton X-100, 0.1% v/v Tween-20, 10 g paraformaldehyde, pH 7.0) for 4 h at 4 °C in the dark. Fixed samples were dehydrated in 30% ethanol for 50 min, then transferred to 50 % ethanol for 50 min, and final stored in 70% ethanol. Tissue sections were then prepared by the In situ Hybridization Core Facility in Academia Sinica (http://abrc.sinica.edu.tw/2010/view/?mid=320&fid=39) as described previously (Chen *et al*., 2016).

### Localizations of YFP fusion protein

To express *Sl*SWEET5b-YFP fusion protein in Arabidopsis protoplasts, the *mYFP* gene was amplified with primers (YFP-F and YFP-R, Supplemental Table S1) and cloned into the vector pRT101 via *BamHI* and *XbaI* sites. The *SlSEET5b* cDNA (714 bp) without the stop codon was amplified using specific primers (SacI-SWT5b-F and BamHI-SWT5b-dTGA-R, Supplemental Table S1), and cloned into the pRT101-YFP via *SacI* and *BamHI* sites (pRT101-SlSWEET5b-YFP). Arabidopsis protoplasts were extracted and transformed with the expression construct as described previously (Ko et al., 2021). To mark the position of inner membranes, the plasma membrane marker *At*PIP2A:CFP, or the vacuolar membrane protein AtγTIP:CFP were co-expressed with *Sl*SWEET5b-YFP in Arabidopsis protoplasts. The YFP fluorescence was visualized by excitation with at 514 nm and emission between 516 and 560 nm. The CFP fluorescence was visualized by excitation with at 458 nm and emission between 456 and 515 nm. The autofluorescence of chloroplast was visualized by excitation with at 561 nm and emission between 562 and 758 nm.

### Yeast growth assay

To express *Sl*SWEET5b in yeast, the cDNA was amplified with specific primers (SWT5b-F and SWT5b-R), cloned into the pDONR221-f1 vector, and then transferred to the pDRf1-GW vector using Gateway technology to produce the pDRf1-SlSWEET5b. The plasmid to express AtSUC2 was construct previously (Ho et. al., 2020). The yeast strain EBY4000 was then transformed with this plasmid for growth assay as described (Ho et al., 2019).

### Transport kinetics in yeast

The radiotracer uptake assay was modified slightly from the previous study (Ho et al., 2019). An overnight cell culture containing the pDRf1-SlSWEET5b plasmid was prepared using fresh synthetic deficient media without uracil (SDM-U media, including 1.7 g yeast nitrogen base without amino acids, 5 g ammonium sulfate, 2% maltose, 2% agar and 0.01% of histidine, leucine and tryptophan in 1 liter). Cells were diluted with fresh media to an OD600 of 0.2 and grown at 30 °C to an OD600 of 0.5. Yeast cells were washed and re-suspended in 50 mM SP buffer (sodium phosphate buffer, containing 112 mg of Na_2_HPO_4_ and 6.8 g of NaH_2_PO_4_ in 1 liter, pH 5) to an OD600 of 5 for the [^14^C]glucose uptake assay as described previously (Ho et al., 2019). Radioactivity of lysed cells was quantified using a Tri-Carb 4810TR scintillation counter (Perkin Elmer, USA) and a kinetic curve determined using single rectangular regression (Sigmaplot Version 13).

### Establishment of RNAi transgenic plants

To make a self-complementary hairpin RNA for RNA interference (RNAi) post-transcriptional silencing, forward and reverse partial coding sequences of *SlSWEET5b* (202-501 bp downstream from ATG) were amplified with specific primers (SWT5b-RNAi-SacI-F and SWT5b-RNAi-SpeI-R for the forward; SWT5b-RNAi-NcoI-F and SWT5b-RNAi-SacII-R for the reverse, Supplemental Table S1). The middle intron loop sequence was amplified from pHellsgate8 plasmid with primers (Intron-F and Intron-R, Supplemental Table S1) and cloned into pRT101 (pRT101-intron). The forward and reverse *SlSWEET5b* partial sequences were then cloned into pRT101-intron via *SacI/SpeI* sites and *NcoI/SacII* sites, respectively. The whole RNAi fragment was then cloned into the binary vector pJH212 via *SacI* and *SacII* sites. The resulting plasmid, pJH212-SlSWEET5b-RNAi, was then transferred into tomato plants by Argrobacterium transformation performed by the Transgenic Plant Core Lab in Academia Sinica (http://abrc.sinica.edu.tw/transplant/index.php). Eighteen T0 transgenic tomato plants were regenerated and transferred to soils. Flowers of 5- to 6-week-old transgenic plants were collected to examine the expression of *SlSWEET5b*. Red fruits of silenced lines were collected to harvest F1 seeds.

### Observation of pollen development

To determine the pollen germination rate, stamens were collected from opened flowers (1 day post anthesis), placed into 1.5 mL Eppendorf tubes, pressed and mixed well with 100 μL pollen germination buffer (20 mM MES, 3mM Ca(NO_3_)_2_, 1 mM KCl, 0.8 mM MgSO_4_, 1.6mM boric acid, 24% w/v PEG4000, 2.5% w/v sucrose, pH6.0) (Lu et al., 2015). The pollen mixture was incubated at 25 °C for 2 h on a shaker at 90 rpm. Pollen germination was deemed successful when the pollen tube was longer than the diameter of a pollen (Cheng et al., 2015). To observe nuclei in pollen, 20 μL of the above pollen mixtures were centrifuged and the supernatant was discarded. Pollens were first fixed for 30 min by adding 50 μL of Carnoy’s fluid (absolute ethyl alcohol : glacial acetic acid = 3 : 1 v/v) (Lu et al., 2015). Subsequently, treated pollen grains were collected by centrifuging at 100xg and stained for 30 min with 20 μL of DAPI (4’,6-diamino-2-phenylindole) solution and observed using a Carl Zeiss LSM780 confocal microscope (Instrument Development Center, NCKU). The DAPI fluorescence was visualized by excitation with at 405 nm and emission between 410 and 492 nm. To observe pollen morphology *in vivo*, stamens were freshly collected and observed on a Cryo Scanning Electron Microscopy performed by Cell Biology Core Lab in Academia Sinica (https://ipmb.sinica.edu.tw/en/core/Plant-Cell-Biology-Core-Lab).

### Sugar content analysis

Stamens and petals were collected from flower buds of 7 mm length and ground into powder with liquid nitrogen. Samples were extracted in 1 mL 80% v/v ethanol at 80°C for 30 min and centrifuged at 13000 x g for 10 min. The supernatant was collected and dried using a Vacufuge Plus concentrator (Eppendorf, NY, USA), and then re-suspended with 50 μL water. To determine contents of glucose, fructose and sucrose, 20 μL of the extract was mixed with 190 μL of the reaction buffer (100 mM pH 7.5 HEPES, 10 mM magnesium chloride, 2 mM adenosine 5′-triphosphate, 0.8 mM NAD, 0.05% v/v Glucose-6-phosphate-dehydrogenase). The NAD(P)H absorption changes at λ = 340 nm in a coupled enzymatic assay were determined by using a microtitre plate reader as described previously (Rodrigues et al., 2020).

### Invertase activity assay

Fresh tomato flower tissues were harvested, immediately freeze dried and ground to powder in liquid nitrogen. Samples were extracted three times with total 1.4 ml extraction buffer and centrifuged at 14000 x g at 4 °C for 10 min. The supernatants were used for activity assay of cytosolic invertase and vacuole invertase and the pellet was for cell wall invertase activity assay as reported previously (Tomlinson et al., 2004) with slight adjustments. In short, 10 μl of supernatant or pellet fractions were added to 180 μl of invertase assay mix. The whole mixtures were incubated at 30 °C for 1 h, then at 85 °C for 3 min. Fructosidase assays were performed by adding 100 μl of the mixture into 400 μl of fructose assay mix. Production of G6P from glucose was determined by the increase absorbance at λ = 340 nm. The extraction buffer, invertase assay mix and fructose assay mix were prepared as described previously (Tomlinson et al. 2004).

## ACKNOWLEDGEMENTS

This work was financially supported by grants from the Ministry of Science and Technology, Taiwan (MOST 105-2628-B-006-001-MY3; MOST 108-2314-B-006-077-MY3) to WJG and by the Australian Research Council (grant no. DP180103834) to YLR. Work in the lab of HEN was financially supported by the German federal state of Rhineland Palatinate within the BioComp project. We also acknowledge the support by Dr. Swee-Suak Rachel Ko, Dr. Kuan and Dr. Wann-Neng Jane in the Core Facility at Academia Sinica (Taipei, Taiwan) to insist tissue sections, tomato transformation and electron microscopy, respectively. We also greatly appreciate constructive comments from Dr. Peter Goldsbrough in Purdue University.

## SUPPLEMENTAL MATERIALS

**Supplemental Table S1. The list of primers used in this study.**

**Supplemental Figure S1. Phenotypes of T0 transgenic RNAi plants.**

**Supplemental Figure S2. Fruit and seed yield of T0 transgenic plants.**

**Supplemental Figure S3. Growth of T1 transgenic plants carrying *SlSWEET5b-RNAi* construct.**

